# Integrated Transcriptomics and Histological Analysis Reveals Gill Adaptation Mechanisms to Hypoxic Stress in Largemouth Bass (*Micropterus salmoides*)

**DOI:** 10.64898/2026.09.10.750788

**Authors:** Xiaoli Ma, Baofeng Su, Anxin Ye, Renyu Wan, Keyan Zhao, Liang Zhong, Wenxin Yu, Yuhe Liu, Jinyi Wu, Wenlong Cai

**Author notes:** Corresponding author: Wenlong CAI, Address: Department of Infectious Diseases and Public Health, Jockey Club College of Veterinary Medicine and Life Sciences, City University of Hong Kong, Kowloon Tong, Hong Kong, China.

## Abstract

Largemouth bass (*Micropterus salmoides*) is a globally significant freshwater aquaculture species known for its rapid growth, high-quality flesh, and excellent nutritional value. Given the increasing incidence of hypoxia in its aquaculture setting, understanding the molecular mechanisms underlying gill adaptation to hypoxic stress is critical for improving resilience in this species. To achieve this, we employed full-length transcript sequencing using PacBio Iso-Seq and high-throughput Illumina RNA-seq to construct a comprehensive transcriptome profile. Histological analyses revealed apparent structural alterations in gill tissues under hypoxic stress (dissolved oxygen: 2.0 ± 0.5 mg/L, duration: 96 h), including curved and swollen gill filaments, apoptosis, and epithelial cell vacuolation. Comparative transcriptomic analysis revealed pronounced shifts in the expression of genes involved in energy metabolism, angiogenesis, immune responses, cell proliferation, and apoptosis. Notably, the down-regulation of genes associated with cell cycle and proliferation suggests that *M. salmoides* may conserve energy by suppressing cell division under hypoxic conditions. In contrast, the increased expression of apoptosis-related genes, including Caspase 7, indicates that apoptosis may play a protective role to maintain tissue homeostasis under hypoxia. Furthermore, the suppression of chemokine-mediated immune pathways implies that hypoxia redirects energy toward essential metabolic processes at the cost of local immune responses. GO and KEGG enrichment analyses identified TNF-α and chemokine-mediated signaling pathways as key regulators for tissue homeostasis and adaptive responses to hypoxic stress. These findings enhance our understanding of the mechanisms underlying hypoxia adaptation and provide valuable genetic resources for breeding strategies aimed at enhancing hypoxia tolerance in *M. salmoides*.

## 1. Introduction

Dissolved oxygen (DO) is a critical environmental variable that shapes fish growth, physiological performance, and survival throughout the life cycle (Zhang et al. 2025). Global climate change, increased aquaculture density, and eutrophication have exacerbated the instability of DO levels in aquatic environments, posing significant challenges to fish health and aquaculture sustainability (Abdel-Tawwab et al. 2019; Qiang et al. 2019). In the process of aquaculture, hypoxia and anoxic environments affect the normal physiological metabolism and a series of life activities of fish species (Shang and Wu 2004). This might lead to suppressed immunity, growth inhibition, and even mass mortality, therefore posing a major threat to the stability of natural populations and aquaculture production.

Among fish organs, the gills play a central role in gas exchange, osmoregulation, and excretion, making them particularly susceptible to hypoxic stress (Abdel-Tawwab et al. 2019; Li et al. 2024). For example, the scaleless carp (*Gymnocypris przewalskii*) and goldfish (*Carassius auratus*) undergo gill remodeling and regulate their ionic state in response to hypoxia (Matey et al. 2008; Mitrovic et al. 2009). The crucian carp (*Carassius carassius*) adapts to hypoxic gills by altering morphology through increased apoptosis and decreased cell proliferation, thereby increasing respiratory surface area (Sollid et al. 2003). The blunt snout bream (*Megalobrama amblycephala*) remodels gill structures by changing lamellar height and area according to oxygen levels (Chen et al. 2017). At the molecular level, oxygen-sensing proteins in fish gill tissues, including *HIFs* (Hypoxia-Inducible Factors), *PHDs* (Prolyl Hydroxylases), and *VHL* (Von Hippel-Lindau protein), served as core regulatory factors in response to low dissolved oxygen (Chen et al. 2017). These proteins are activated to varying degrees in hypoxic environments, as observed in species such as the mudskipper (*Boleophthalmus pectinirostris*) (Xiao 2015a), silver sillago (*Sillago sihama*) (Saetan et al. 2020), and channel catfish (*Ictalurus punctatus*) (Geng et al. 2014). Under normoxic conditions, mitochondria in fish produce a small amount of reactive oxygen species (ROS) to support normal physiological activities (Kulisz et al. 2002). However, under hypoxic stress, the dynamic equilibrium of ROS is disrupted, leading to excessive production of ROS, which subsequently causes oxidative damage in fish. For instance, in juvenile Japanese seabass (*Lateolabrax japonicus*), oxidative stress indicators and antioxidant enzyme activities in the gill tissues were significantly upregulated under hypoxic conditions (Peng 2020). All these changes reflect the adaptive effects to hypoxic stress, enabling better survival in low oxygen environments.

To date, many fish species have been shown to cope with or adapt to hypoxia through complex molecular strategies. Hypoxia in Japanese flounder (*Paralichthys olivaceus*) triggers the activation of the HIF- 1/LDH-A and EPAS1/Bad signaling pathways, influencing gene expression and DNA methylation, thereby inducing physiological alterations and apoptosis in the gills (Liu et al. 2022; Li et al. 2022). Changes in the expression of genes involved in anaerobic glycolysis, fatty acid β-oxidation, glutathione metabolism, peroxisomal antioxidant functions, and apoptosis play important roles in the transcriptome response to hypoxia in the pearl gentian grouper (*Epinephelus lanceolatus*♂×*Epinephelus fuscoguttatus*♀) (Jiang et al. 2023). Hypoxic stress leads to significant transcriptional regulation related to glycolipid metabolism in Hulong hybrid grouper (*Epinephelus fuscoguttatus*♀×*E. lanceolatus*♂), in which insulin, HIF-1, FoxO, and cAMP pathways play key roles (Wang et al. 2024). The blunt snout bream (*Megalobrama amblycephala*) displays numerous differentially expressed genes (DEGs) after hypoxic stress. These genes were involved in multiple biological processes, such as the HIF-1, FOXO, MAPK, PI3K-Akt, and AMPK signaling pathways (Zhao et al. 2022). Upregulation of apoptosis-related genes and enhanced antioxidant capacity under hypoxic conditions have also been observed in *Pelteobagrus vachelli* (Zheng et al. 2021). Additionally, hypoxic stress leads to immune responses in various fish. Hypoxia modulates innate immune activities, including cell migration, programmed cell death, pathogen phagocytosis, antigen processing, as well as the release of cytokines, chemokines, angiogenic mediators, and antimicrobial peptides (Sarah et al. 2014). In Large yellow croaker (*Larimichthys crocea*), hypoxia significantly downregulates a variety of pattern recognition receptors (PRRs), including Toll-like receptors (*TLR1*, *TLR2-1*, *TLR2-2*, *TLR5* and *TLR8*), fucolectins (*FUCL1*, *FUCL4* and *FUCL5*), and macrophage mannose receptor (*MRC1*), suggesting that hypoxic stress inhibits immune processes mediated by these PRRs (Mu, Li, Wu, et al. 2020). The Nod-like receptor-mediated immune response plays an important role in hypoxia tolerance in hybrid yellow catfish (*Tachysurus fulvidraco*♀×*Pseudobagrus vachellii*♂)(Qiang et al. 2021). It was also reported that chronic hypoxia alters the expression of eight key immune genes (*IFN*α, *IFN-*γ, *Mx*, *MxA*, *MxB*, *IL-1*β, *TNF-*α*1*, and *IP10*) in Atlantic salmon (Kvamme et al. 2013).

*M. salmoides* is a fast-growing, temperature-tolerant fish with a high degree of plasticity in adapting to new environments. Its rapid growth, ease of farming, and short breeding cycle make it one of the major cultured fish species in China (Bai et al. 2008). Moreover, *M. salmoides* is highly valued by farmers for its ability to thrive in various aquaculture environments (Masagounder et al. 2009). However, the thriving *M. salmoides* industry faces significant challenges from environmental problems such as hypoxia, which severely restrict the stable, healthy and sustainable development of its aquaculture. Despite its economic importance, few studies have investigated the molecular mechanisms of *M. salmoides* under low oxygen stress. In this study, we integrated PacBio full-length transcriptome sequencing, Illumina RNA-seq, and histological analysis to systematically investigate gill responses of *M. salmoides* to hypoxic stress. Full-length transcriptome sequencing provided improved transcript structural annotation and isoform-level information, whereas comparative RNA-seq and histological analyses revealed hypoxia-induced changes in gene expression and gill morphology. Together, these analyses identified key molecular pathways and tissue-level features associated with hypoxia adaptation, providing valuable resources and insights for future studies on hypoxia tolerance and selective breeding in *M. salmoides*.

## 2. Materials and methods

### 2.1 Ethical statement

This study complied with the "Guidelines for the Welfare and Ethical Review of Experimental Animals (GB/T 35892-2018)." All procedures involving animal handling and tissue collection were approved by the Institutional Animal Care and Use Committee at Jiangsu Normal University. Prior to sampling, fish were anesthetized using MS-222 (tricaine methanesulfonate) to minimize suffering.

### 2.2 Hypoxia challenge and sample collections

All fish used in this experiment (body length: 15.0 ± 1 cm, weight: 35 ± 5 g) were obtained from Fishery Life Aquatic Product CO., Ltd. (Wuxi, Jiangsu, China). Ninety fish were randomly transferred to six glass aquaculture tanks (48 cm × 30 cm × 25 cm) equipped with biofilters for 7-day acclimation, with 15 fish per tank. The experimental water temperature was maintained at 19.5 ± 1□, with the dissolved oxygen (DO) maintained at 7.3 ± 0.5 mg/L, and pH at 8.1 ± 0.2. The fish were fed twice daily with artificial compound feed at 9:00 a.m. and 5:00 p.m. Feed was withheld for 24 h before the start of the experiment.

Fish were assigned to a hypoxia treatment group (EG, DO 2.0 ± 0.5 mg/L) or a normoxic control group (CG, DO 7.0 ± 0.5 mg/L), with three replicate tanks per group. During the hypoxia challenge, the aeration equipment for the EG group was turned off, and pure nitrogen was bubbled into the water for approximately 30 min to lower the DO concentration to 2.0 mg/L. Subsequently, the nitrogen influx was adjusted to maintain the DO concentration at 2.0 ± 0.5 mg/L. The DO concentration in the water was measured every 30 min throughout the experiment using a dissolved oxygen meter (AR8046, SMART SENSOR, Guangdong, China).

After 96 h, the experiment was terminated. One fish was randomly selected from each tank, anesthetized with MS-222 (300 mg/L), and its gill tissue was collected. The gill tissue was then treated with liquid nitrogen and stored at -80 for further analysis. Additionally, another sample of gill tissue was collected and stored in a centrifuge tube containing 4% paraformaldehyde for histological sectioning and examination.

### 2.3 RNA extraction, library construction and sequencing

Total RNA was extracted using the Trizol reagent kit (Invitrogen, Carlsbad, CA, USA) according to the manufacturer’s protocol (Cai et al. 2022). RNA integrity was evaluated using an Agilent 2100 Bioanalyzer and agarose gel electrophoresis. RNA purity and concentration were determined using a Nanodrop micro- spectrophotometer (Thermo Fisher Scientific). mRNA was enriched from total RNA using Oligo(dT) beads provided with the Hieff NGS® mRNA Isolation Master Kit. For PacBio Iso-Seq, the enriched mRNA was reverse-transcribed into cDNA using the Clontech SMARTer PCR cDNA Synthesis Kit. PCR cycle optimization was conducted to determine the optimal number of cycles for large□scale downstream PCR amplification to generate double stranded cDNA. Size selection (> 5 kb) was performed using the BluePippin™ Size Selection System, and the size□selected fraction was mixed equally with the non size selected cDNA. Large□scale PCR was then performed for subsequent SMRTbell library construction. The resulting cDNA products were subjected to DNA damage repair, end repair, and adapter ligation to prepare SMRTbell templates. The SMRTbell templates were annealed to sequencing primers, bound to polymerase, and sequenced on the PacBio Sequel II platform by Gene Denovo Biotechnology Co. (Guangzhou, China). The raw data were deposited in the National Center of Biotechnology Information (NCBI) database under accession number PRJNA1501902.

For Illumina RNA-Seq, mRNA was fragmented and reverse transcribed into cDNA using the NEBNext Ultra RNA Library Prep Kit for Illumina (NEB #7530, New England Biolabs, Ipswich, MA, USA). The purified double-stranded cDNA was end-repaired, A-tailed, and added to Illumina sequencing adapters. The ligation reaction was purified with the AMPure XP Beads (1.0X) and PCR amplified. All libraries were constructed in three bio-logical replicates for each group, resulting in a total of six cDNA libraries. The resulting cDNA libraries were sequenced using the Illumina NovaSeq 6000 by Gene Denovo Biotechnology Co. (Guangzhou, China). The raw reads were deposited in the NCBI database under accession number PRJNA1498806.

### 2.4 Data analysis for Iso-Seq

Raw cDNA sequencing reads were processed using the Iso-Seq workflow supported by Pacific Biosciences (Gordon et al. 2015; Zhang et al. 2024). First, high-quality circular consensus sequences (CCS), also called HiFi reads, were extracted from the subreads BAM file. The integrity of transcripts was assessed by checking whether the CCS reads contained 5’ primer, 3’primer, and poly(A) structures. Sequences containing all three structures were considered full-length sequences (FL reads).

Afterwards, primers, barcodes, poly(A) tails, and concatemer sequences were removed from full-length reads to obtain full-length non-chimeric (FLNC) reads. The FLNC reads were then clustered using the ISO-Seq clustering pipeline to generate unpolished consensus isoforms (Li 2018). The Quiver algorithm was used to further correct the consensus sequence. High-quality isoforms with a prediction accuracy ≥ 0.99 were selected for the subsequent analysis. Using Minimap2, the corrected and merged sequences were mapped to the reference genome (GenBank Accession No. GCA_036785525.1), and the clustering output was used to derive nonredundant transcripts.

To investigate the functions of new isoforms, new isoforms were BLAST against the NCBI non-redundant protein (Nr) database, the Swiss-Prot protein database, and the Kyoto Encyclopedia of Genes and Genomes (KEGG) database using the BLASTx program (http://www.ncbi.nlm.nih.gov/BLAST/) at an E-value threshold of 1e-5 (Gao et al. 2024). Gene Ontology (GO) annotation was performed using Blast2GO software based on Nr annotation results (Conesa et al. 2005).

Two software tools, CNCI (version 2) and CPC (http://cpc.cbi.pku.edu.cn/), were used to evaluate the coding potential of transcripts under their default settings (Liang et al. 2013; Kong et al. 2007). In parallel, isoforms were searched against the SwissProt database for protein annotation. Transcripts lacking both protein-coding potential and protein annotation were classified as long non-coding RNAs (lncRNAs). Alternative splicing events were identified using the SUPPA (v2.3) tool (Alamancos et al. 2015). Polyadenylation patterns for expressed genes and their transcripts were characterized with Tapis (v1.2). The SAMtools (v1.10) mpileup coupled with BCFtools (v1.18) call were used for calling variants of transcripts with thresholds of mapping quality ≥ 20 and base quality ≥ 20, and ANNOVAR was used for SNP/InDel annotation (Li et al. 2009; Wang, Li, and Hakonarson 2010). The function, genomic location, and type of variation of SNPs were also analyzed. Protein coding sequences were scanned against AnimalTFDB using HMMER (v3.3.1) hmmscan to identify transcription factor (TF) families (Huang et al. 2025).

### 2.5 Data analysis for Illumina RNA-Seq

The raw reads were filtered by fastp (version 0.18.0) to remove the adaptor and low-quality reads (Chen et al. 2018). The paired-end clean reads were mapped to the reference genome (GCA_036785525.1) using HISAT2 (v2.1.0), with other parameters set as default (Kim, Langmead, and Salzberg 2015). Mapped reads from each sample were assembled with StringTie (v1.3.1) in a reference-guided approach, and transcript abundance was estimated as FPKM values using RSEM (v1.2.19) (Pertea et al. 2015; Kim et al. 2016; Dewey and Bo 2011).

Differential expression analysis was carried out using DESeq2 (v1.24) (Love, Huber, and Anders 2014). Genes with a false discovery rate (FDR)-corrected *P*-value < 0.05 (Benjamini–Hochberg correction) and |log2(FC)| > 1 were defined as differentially expressed genes (DEGs). Principal component analysis (PCA) was performed using the R package gmodels in this research. The DEGs were visually presented in a Volcano Plot and Heatmap. To gain insight into the molecular mechanisms and biological processes associated with genes involved in the hypoxic response, the DEGs were mapped to Gene Ontology (GO) and Kyoto Encyclopedia of Genes and Genomes (KEGG) databases for enrichment analysis. The clusterProfiler (v4.2.0) (Yu et al. 2012) was used to conduct enrichment analyses of DEGs, with the Benjamini–Hochberg procedure for multiple-testing correction and an adjusted *P*-value (FDR) < 0.05 as the significance threshold.

### 2.6 Histological analysis

At the end of the exposure, three fish were randomly selected from each treatment group for gill tissue collection. Each sample was separately placed in a 4% paraformaldehyde solution for 24 h of fixation and then sent to Servicebio Technology (Wuhan, China) for Hematoxylin and Eosin (HE) staining. Gradient dehydration was performed with alcohol, then xylene was used for tissue clearing, followed by wax dipping and embedding. After embedding, tissue sections (4 μm in thickness) were prepared and fixed on slides. Sections were then deparaffinized, stained with hematoxylin-eosin, dehydrated, sealed, and examined microscopically for image acquisition and analysis. For morphometric analysis, five gill filaments were randomly selected from each histological image and their measurements were averaged. The length (red arrows) and thickness (black arrows) of these selected gill filaments were quantified using Image-Pro Plus 6.0 software (Media Cybernetics, U.S.A), with millimeters (mm) as the standard unit.

## 3. Results

### 3.1 Full-length transcriptome construction and annotation

There were no mortality cases in either group after 96 h. *M. salmoides* in the hypoxic group exhibited slower swimming and reduced feeding behavior. To investigate the gene expression patterns in gills under hypoxic stress, we collected gill samples for full-length transcriptome and RNA-seq analyses. The full-length transcriptome sequencing produced 31,591,017 subreads, with an average length of 2,589 bp and an N50 value of 2,638 bp (Fig. S1a). A total of 489,744 CCS sequences were identified after filtering. The number of FLNC reads was 419,466, accounting for 85.65% of the CCS reads, with an average length of 2,580 bp (Fig. S1b). After clustering and further correction of the unpolished consensus isoforms using the Quiver algorithm, we obtained 27,731 high-quality (HQ) isoforms and 191 low-quality (LQ) isoforms. All high-quality (HQ) isoforms were aligned to the reference genome using Minimap2, resulting in 27,377 aligned isoforms (98.72%) and 354 unaligned isoforms (1.28%) (Fig. S1c). We retained the mapped isoforms and removed redundant transcripts to generate a nonredundant transcript set. This nonredundant collection (24,464 mapped isoforms) was subjected to genome alignment again for isoform classification. Among these 24,464 mapped isoforms, 14,056 (57.5%) matched previously annotated isoforms (Table S1). Isoforms that align to unannotated regions of the genome are considered as novel isoforms, while those that align to consistent sequences of different exons from known isoforms are regarded as new isoforms. A total of 554 novel isoforms and 9,854 new isoforms were annotated using the NR, GO, KEGG, and SwissProt databases together with the reference-based gene annotation, with 10,070,111,050,1,487 and 9,037 acquired annotations.

GO annotation analysis assigned these isoforms to biological processes such as metabolic regulation, immune response, and response to stimulus (Fig. S1d). KEGG pathway analysis revealed significant enrichment in metabolism and organismal systems related pathways. In particular, metabolic pathways were associated with 1,453 genes, followed by complement and coagulation cascades with 498 genes (Table S2).

### 3.2 Gene structure analysis of Iso-Seq

Integration of lncRNA predictions from three independent tools identified 442 lncRNAs, which were retained as the final non-coding RNA set (Fig. S2a). LncRNAs were classified into 5 types based on their genomic positions relative to protein-coding genes: intergenic, intronic, sense, antisense, and bidirectional lncRNAs (Fig. 1a). Alternative splicing is important for regulating gene expression and generating protein diversity. A total of 4,120 AS events were identified, which were classified into seven types of AS events. Among them, retained introns (RI) were the most prevalent, accounting for 1,021 events (Fig. 1b). A comparative analysis of the two groups revealed a significant induction of variable shear changes in genes, with a total of 522 significantly different variable shear events identified. Among these, exon skipping (SE) was the most predominantly affected type (397 events), and exon skipping was slightly facilitated (i.e., the inclusion level was reduced) by hypoxia treatment. Changes in the variable 3’ shear site (A3SS) were also significant (59 events) and tended to increase the inclusion level of the variable region. Using Tapis software, 24,464 polyA sites were identified from 11,569 genes (Fig. S2b).

**Fig. 1.**
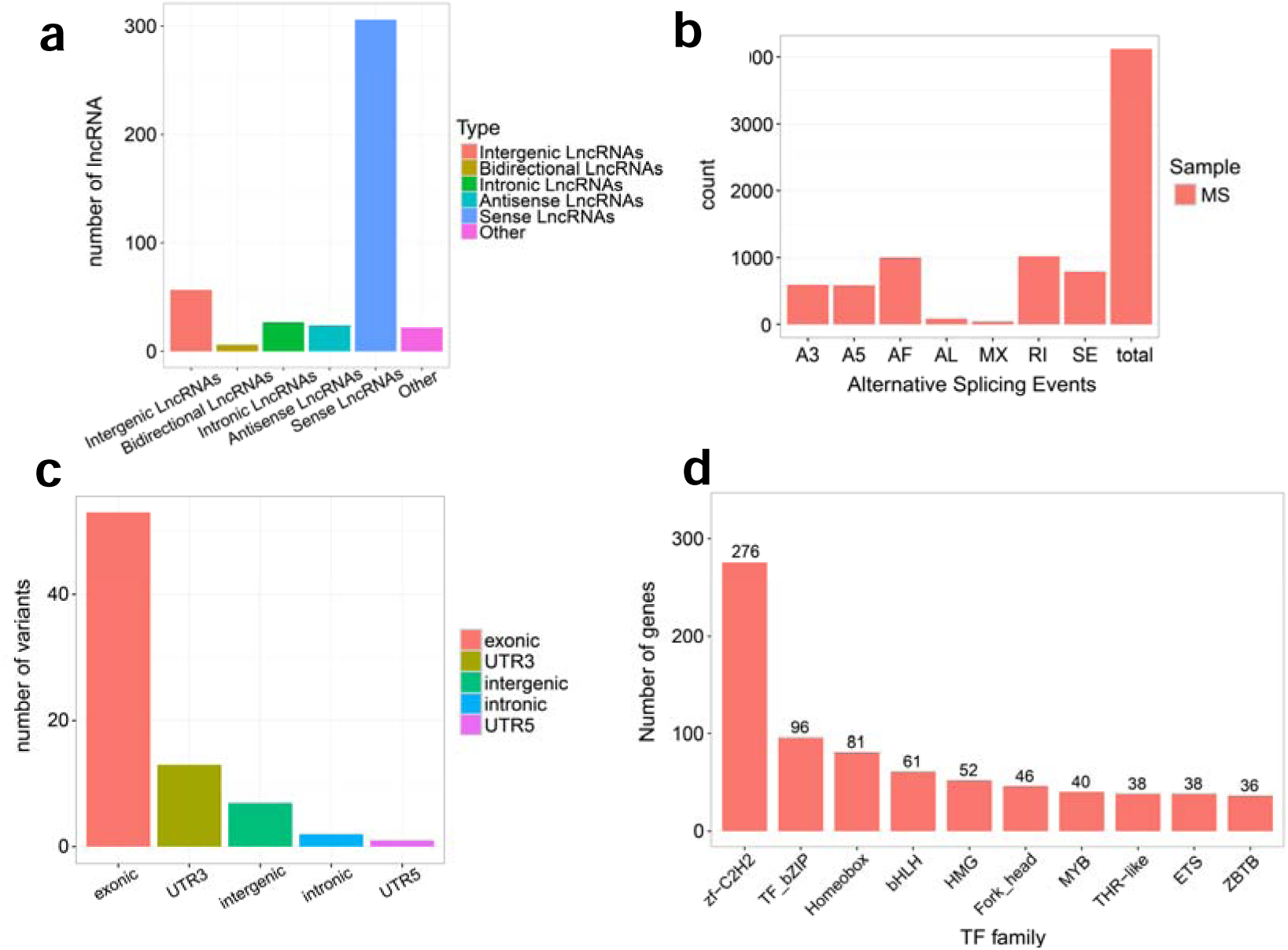
(a) Summary of lncRNA classification results. (b) Summary of alternative splicing (AS) events. A3, alternative 3′ splice site; A5, alternative 5′ splice site; AF, alternative first exon; AL, alternative last exon; MX, mutually exclusive exon; RI, retained intron; SE, skipped exon. (c) Classification of SNP positions. The X-axis represents different location types, and the Y-axis represents the number of variants. (d) Results of distribution of top 10 transcription factor (TF) families.

A total of 53 SNPs were identified, including 26 synonymous and 24 nonsynonymous variants (Fig. S2c). The classification map of location types produced by SNPs indicates five distinct categories. The statistical analysis of SNP mutation types showed that transition mutations, particularly A-> G substitutions, represent the predominant SNP class, while transversions occurred less frequently. Among transversions, C-> G and T-> G were the least common, consistent with the general observation that transitions outnumber transversions in genomes (Fig. S2d). The gene variations are primarily concentrated in the exonic regions. The number of variants in the UTR3 and intergenic regions is relatively higher, while the intronic and UTR5 regions exhibit notably fewer variants (Fig. 1c). This distribution pattern provides crucial data for subsequent gene research and variation analysis. Transcription factors (TFs) are a unique set of sequence-specific DNA-binding proteins that regulate the transcription of target genes. Hmmsearch predicted a total of 62 transcription factor families, including 1,172 transcription factors in *M. salmoides*. The zf-C2H2 family had the largest family of transcription factor families, containing 276 members, followed by TF_bZIP (96), Homeobox (81), bHLH (61), HMG (52), Fork_head (46), MYB (40), THR-like (38), ETS (38), and ZBTB (36) (Fig. 1d).

### 3.3 Illumina Sequencing and Reads Mapping

A total of 231,620,654 raw reads were obtained using the Illumina NovaSeq 6000 sequencing platform. Following the removal of low-quality raw reads, 230,660,282 clean reads were obtained. Q20 and Q30 scores were greater than 97% and 92%, respectively, with an average GC content of 47.19% (Table S3). The high-quality reads obtained were aligned with the reference genome (GCA_036785525.1) of *M. salmoides*. The results demonstrated that the total mapping rate and unique mapping rate were greater than 95% and 87%, respectively. Mapped reads were predominantly localized to exons (87.89-88.61%) based on the structure of the reference genome.

### 3.4 Analysis of differentially expressed genes (DEGs)

To identify DEGs, we analyzed gill transcriptome data from *M. salmoides* hypoxic and normoxic groups. The reads count data obtained from RSEM were loaded into the R language package DESeq2 for analysis, which identified 614 DEGs in the treated group (|log2(FC)|>1 and FDR < 0.05), including 372 down- regulated genes and 242 up-regulated genes (Fig. 2a, b; table S4). The application of heatmap and principal component analysis (PCA) revealed that DEGs were clustered into two principal branches, with genes within the same branch exhibiting similar expression patterns under different experimental conditions (Fig. 2c, Fig. 2d). This suggests that hypoxic stress had a significant impact on the gill tissues of *M. salmoides*.

**Fig. 2.**
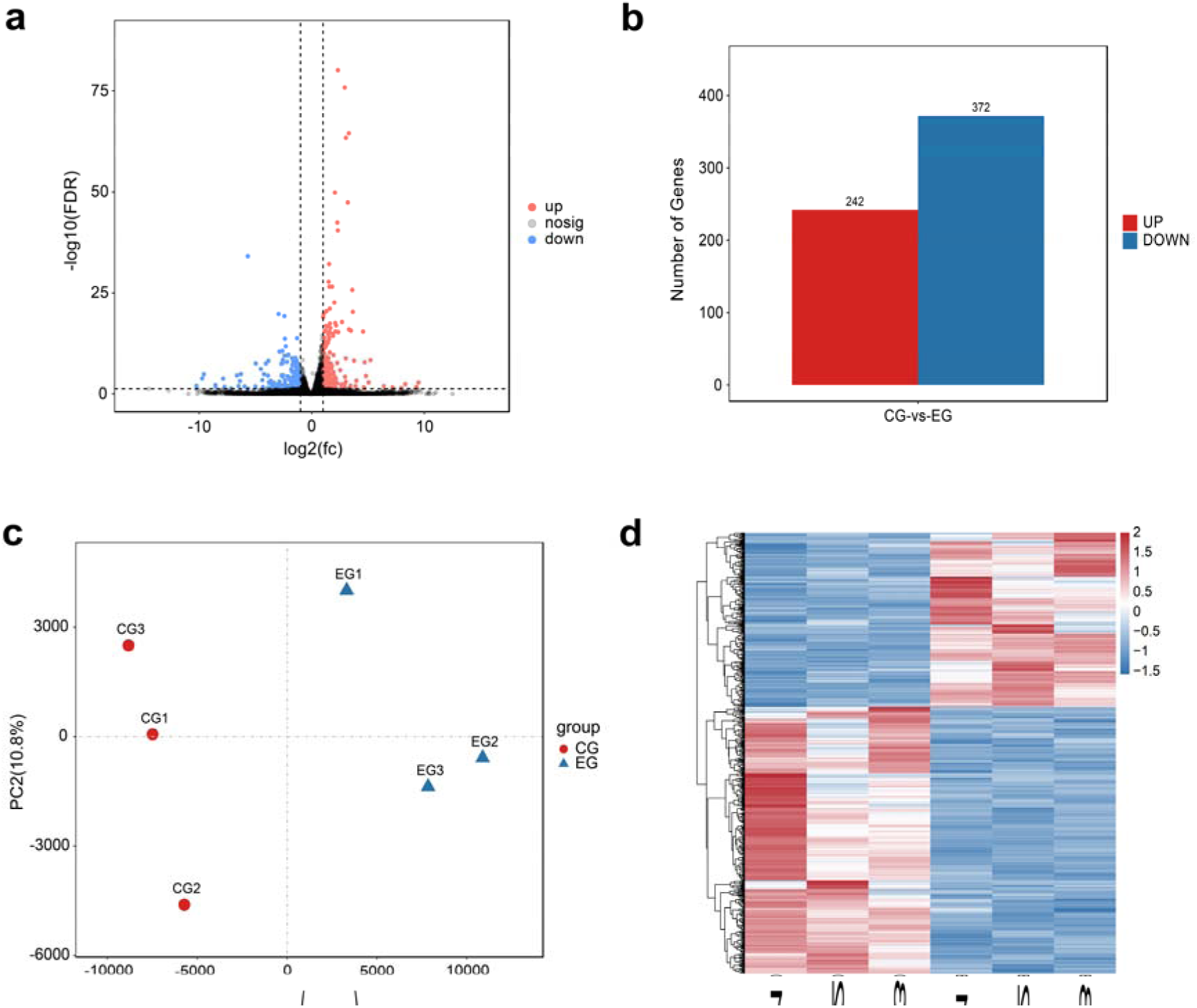
(a) Volcano plot of differentially expressed genes (DEGs). DEGs with |log2(FC)|>1 and FDR < 0.05 were chosen for volcano plot in both groups. Red points indicate the up-regulated DEGs, and blue points indicate down-regulated DEGs. (b) Statistical distribution of DEGs. The X-axis represents the comparison between the control group (CG) and the experimental group (EG), while the Y-axis represents the number of DEGs. Red bars represent up-regulated genes (242), blue bars represent down-regulated genes (372). (c) Principal component analysis (PCA) plot. The X-axis (PC1) explains 84.3% of the total variance, and the Y- axis (PC2) explains 10.8% of the total variance. The red dots represent samples from the control group (CG), and the blue dots represent samples from the experimental group (EG). (d) Heatmap of DEGs. Rows represent DEGs, and columns represent different samples. The color gradient indicates the expression levels, with red indicating up-regulation and blue indicating down-regulation.

### 3.5 GO and KEGG enrichment analysis of DEGs

GO annotation of the 614 DEGs from hypoxic gill tissues showed that most were classified under biological processes (BP, 75.15%), followed by molecular function (MF, 15.09%) and cellular component (CC, 9.76%) (Table S5). In the BP category, 15 hypoxia-related terms were significantly enriched (*P*-value < 0.05), including response to hypoxia, ATP synthesis and ATPase activity, and angiogenesis-related pathways, indicating that *M.salmoides* actively senses hypoxic signals and adapts by modulating oxygen-related response pathways (Fig. 3). Moreover, significant enrichment of molecular function categories such as “CCR chemokine receptor binding”, indicates that hypoxic stress may effect the immune response in gills of *M. salmoides* (Table S5).

**Fig 3.**
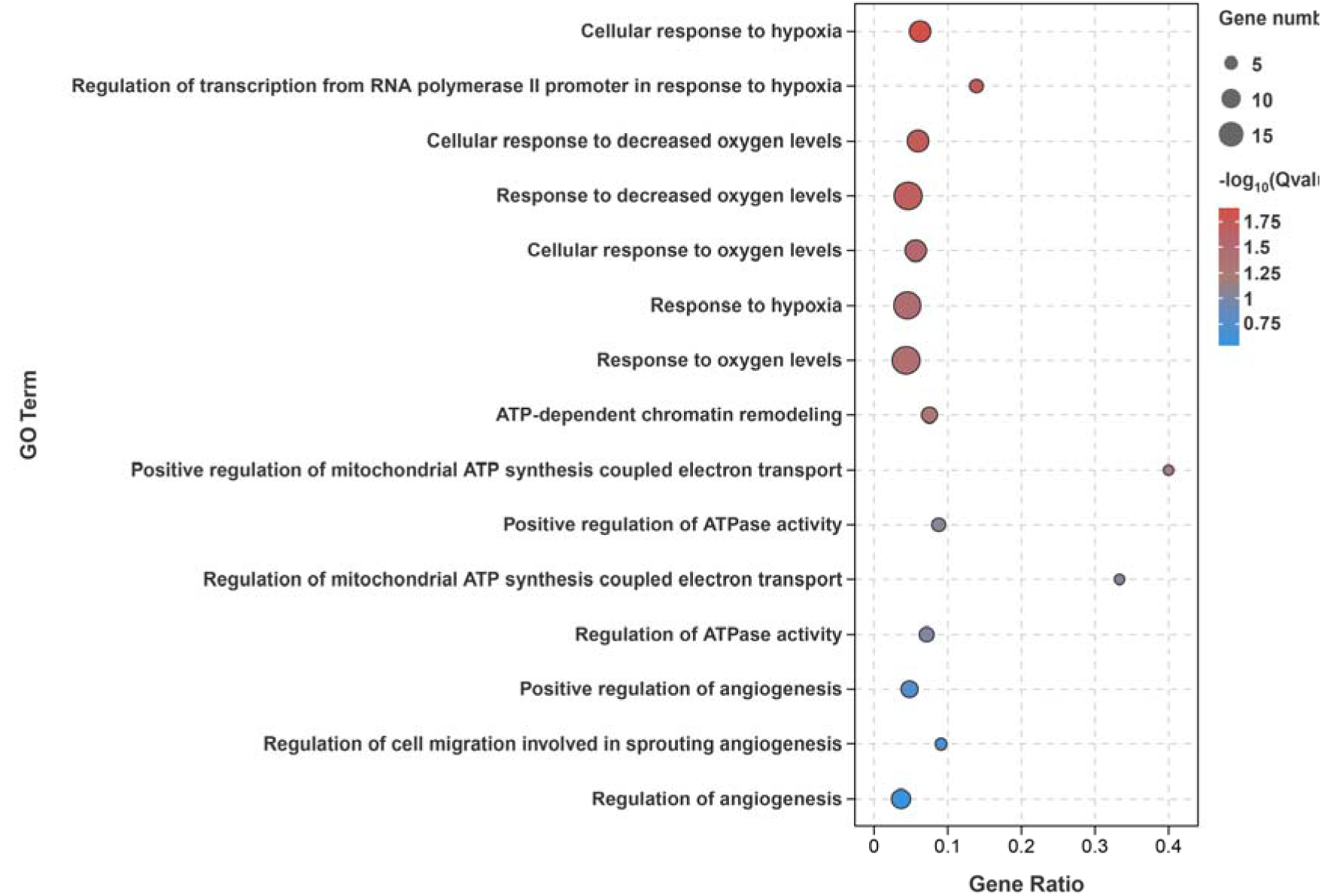
GO enrichment results (BP) of DEGs

In summary, *M. salmoides* demonstrated complex and diverse adaptive strategies by regulating transcriptional programs associated with hypoxia response, angiogenesis, and energy metabolism under hypoxic conditions. These regulatory mechanisms may help the fish to maintain basic physiological functions and ensure its survival in hypoxic environments.

Among the 614 DEGs, 176 were assigned to KEGG pathways, and 27 pathways were significantly enriched (P < 0.05; Table S6). The 20 most significantly enriched pathways ranked by ascending adjusted p-values showed that “DNA replication”, “Cell cycle”, “Cytokine-cytokine receptor interaction”, “Pyrimidine metabolism”, “Toll-like receptor signaling pathway”, “Apoptosis”, and “TNF signaling pathway” were the most abundant pathways enriched in DEGs (Fig. 4).

**Fig 4.**
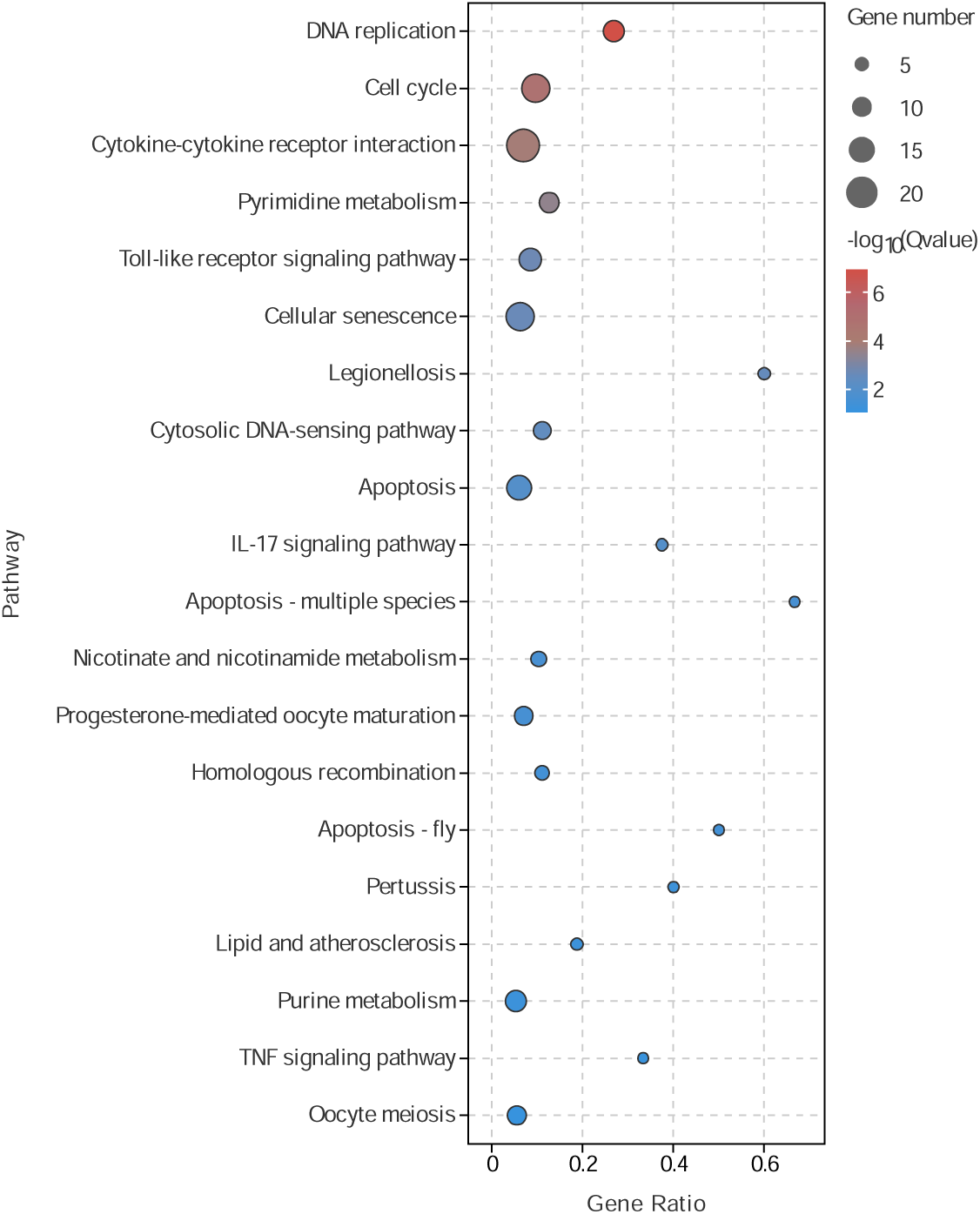
The top 20 enriched KEGG pathways ranked by ascending adjusted P-values

Taken together, the results suggested that the adaptive strategy of *M. salmoides* in a hypoxic environment involves the regulation of cell proliferation, cell cycle, immune response, and cell apoptosis, and these mechanisms together enable *M. salmoides* to survive under low oxygen conditions.

### 3.6 Histological observation of the gill tissue

Histological analysis showed that the gill tissue retained normal physiological morphology under normoxic conditions. The branchial lamella were arranged neatly, the red blood cells were uniformly distributed, the pavement cells were regularly arranged, and the mitochondria-rich cells were distributed in the filament core (Fig. 5a).The gill lamellae on the gill filaments of the *M. salmoides* were short, thick, and symmetrically distributed on both sides of the gill filaments with closely and regularly arranged gill lamellae (Fig. 5c). The gill filaments had a blood sinus in the center, which appears as a long, narrow space with uniformly distributed blood cells (Fig. 5e).

**Fig. 5.**
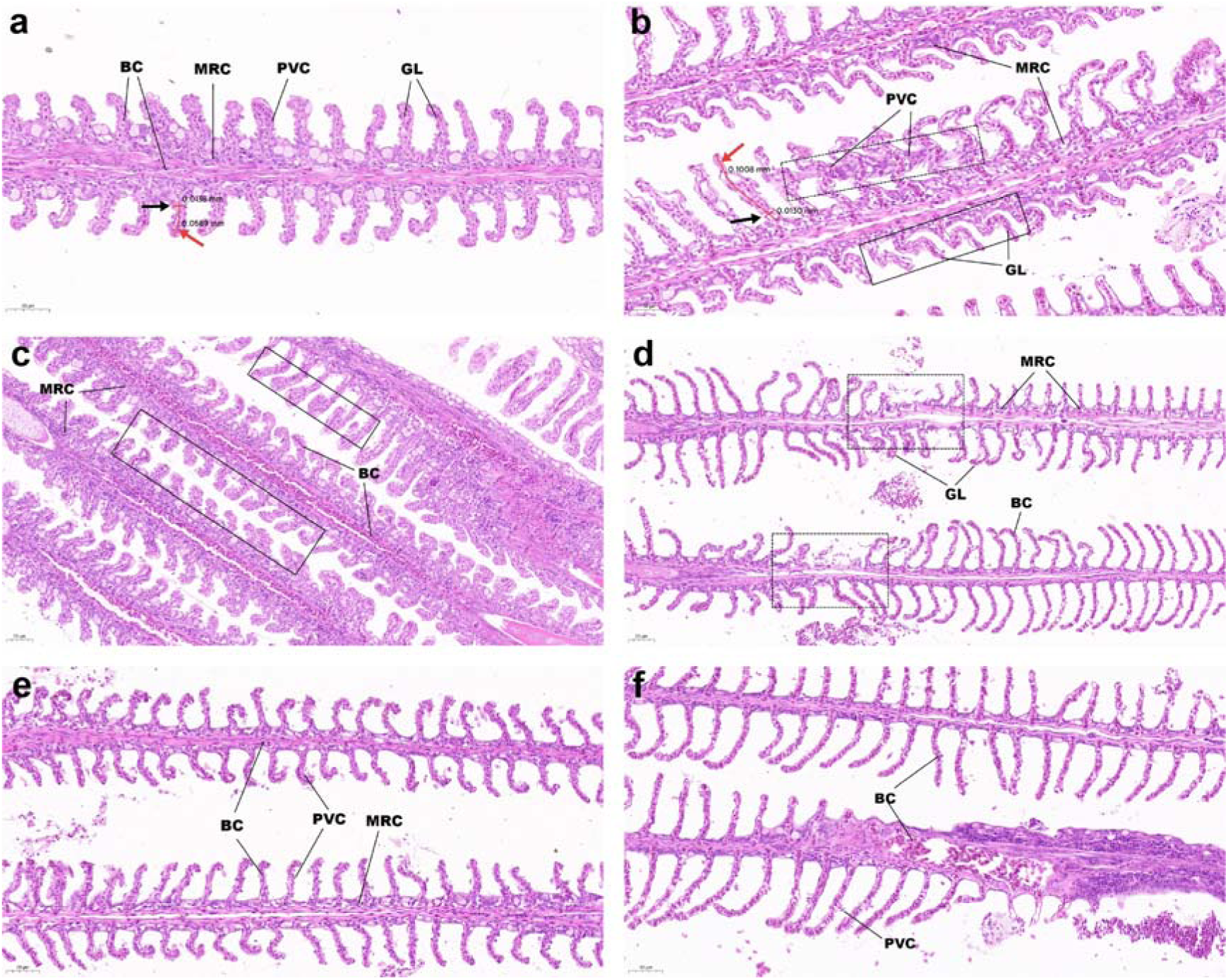
Hematoxylin and eosin (H&E) staining of gill sections under different oxygen conditions: (a)(c)(e) normoxia; (b)(d)(f) hypoxia. GL: gill lamellae ; PVC: pavement cells ; MRC: mitochondria-rich cells ; BC: blood cells

After 96 h of hypoxic stress, mitochondria-rich cells showed an increase in size and number, the gill lamellae adhered to each other, forming an “S” shape, and the pavement cells were swollen and vacuolated, eventually rupturing and shedding into the interstitial space of the lamellae, showing signs of necrosis (Fig. 5b). These changes may increase the gill respiratory surface to enhance oxygen uptake, consistent with the structural changes observed in silver carp under hypoxia (Feng et al. 2022). The gill lamellae became disorganized, showing asymmetry and breakage (Fig. 5d). The sinusoidal space was reduced and the blood cells showed uneven distribution (Fig. 5f). The above results indicate that hypoxia induced significant structural changes in the gills, deviating from the normal gill physiological functions, and may impair the normal gill physiological function. The average gill filament length was 0.0589 mm and 0.1008 mm in the normoxic group and hypoxic group, respectively. In addition, the average gill filament thickness was 0.0138 mm in the normoxic group and 0.0130 mm in the hypoxic group (Table S7). However, these differences were not statistically significant (p > 0.05).

### 3.7 Potential regulatory mechanisms of gill responses to hypoxia

Based on KEGG signaling pathway enrichment analysis, we further deciphered a putative molecular mechanisms underlying gill responses to hypoxic stress in *M. salmoides* (Fig. 6). When exposed to hypoxia, gill cells responded to low-oxygen environments through the upregulation of hypoxia-inducible factor 1-alpha (*hif1*α), which is a key regulator of the hypoxia response. HIF-1α transmitted downstream signals by modulating the expression of vascular endothelial growth factor A (*vegfa*), thereby regulating angiogenesis. VEGFA further propagated signaling through calcium signaling pathway. Meanwhile, gill cells also perceived this hypoxic stress through the calcium signaling pathway, which induced the upregulation of calmodulin (*calm*) gene expression. This upregulated calm subsequently transduced signals to the nucleus, leading to reduced expression of tumor necrosis factor α (*tnf*α) to mitigate cellular inflammatory responses in the gill tissue. Additionally, a low level of *tnf*α was involved in regulating apoptosis. Concurrently, the upregulation of calpain gene expression transmitted pro-apoptotic signals to caspases, resulting in increased caspase 7 expression, which in turn promoted apoptosis in gill cells. Notably, we observed a decrease in the expression of extracellularly secreted substances like *perforin*, which diminished the cytotoxic effects of cytotoxic T lymphocytes (CTLs) and natural killer (NK) cells on target cells, as well as *GZMB*, which was also involved in apoptosis regulation. In addition, transcriptomic and KEGG pathway analyses revealed reduced expression of *CHK1*, a checkpoint-related gene. This reduced *CHK1* then transduced signals to downstream genes via the p53 signaling pathway, leading to downregulated expression of cell cycle-related genes such as *Cdk1*, *Cyclin B (CycB)*, *Cyclin A* (*CycA*), and *E2F* transcription factor family members. These changes ultimately resulted in cell cycle arrest in gill tissues. Furthermore, DNA replication-related genes, such as *MCM*, α*1*, *RPA3*, *Dna2*, and *Fen1*, also showed significant reductions, which contributed to impaired cell proliferation. In summary, we found that hypoxic stress could induce apoptosis and inhibit the cell cycle in the gills of *M*.

**Fig. 6.**
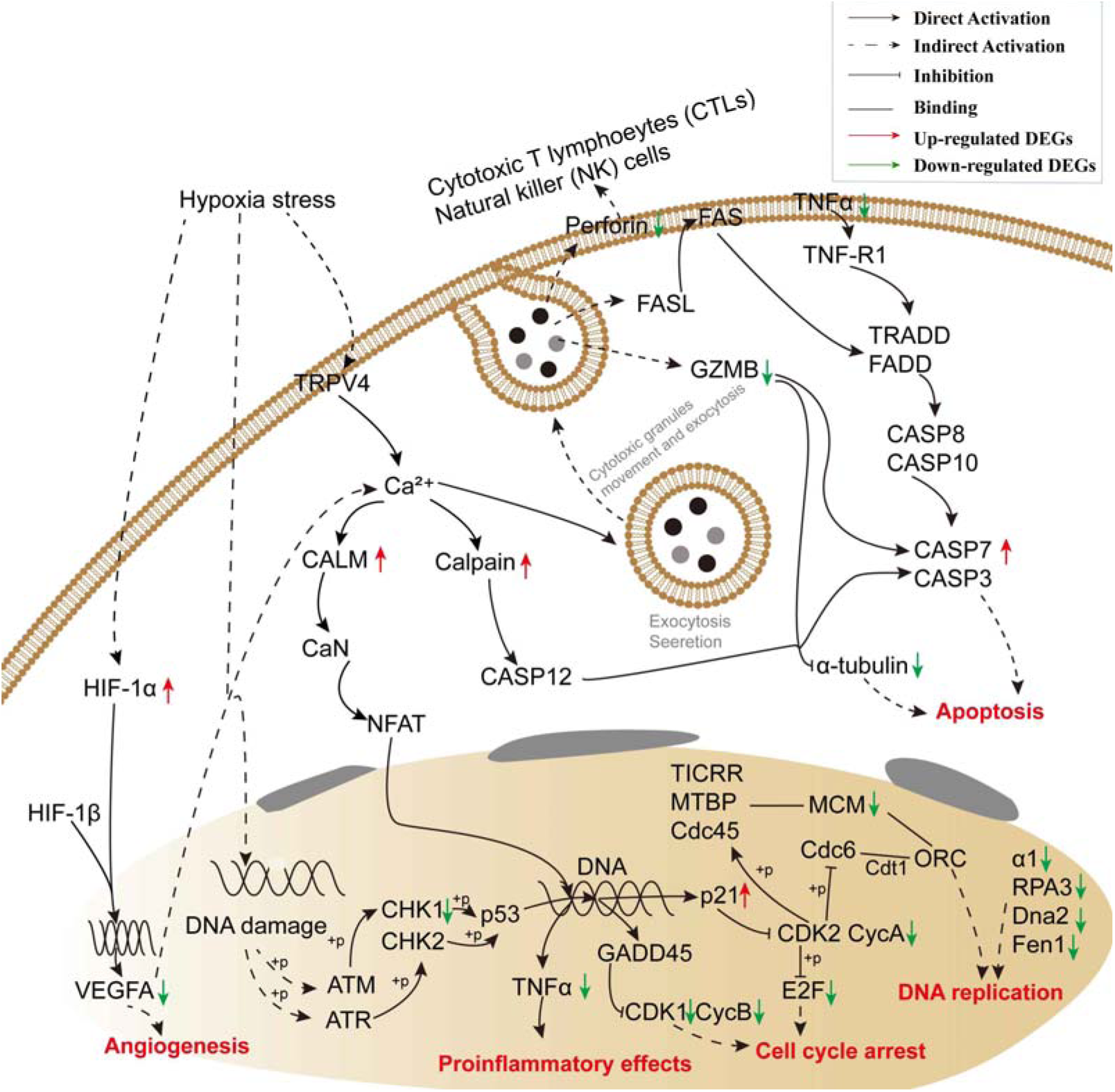
Potential molecular regulatory network underlying *M. salmoides* gill responses to hypoxia, which was mapped based on KEGG signaling pathway enrichment. The molecular interactions, including direct activation, indirect activation, inhibition, and binding, were retrieved from the KEGG database. Red arrows represent significantly up-regulated differentially expressed genes (DEGs), and green arrows indicate significantly down-regulated DEGs identified in our transcriptome data.

## 4. Discussion

### 4.1 Hypoxia induced AS event and transcript variants

This study utilized PacBio Iso-Seq to analyze the full-length transcriptome of *M. salmoides* gills under hypoxic stress. Unlike conventional short-read sequencing, Iso-Seq directly captures complete transcript sequences, which yields more accurate isoform discovery and circumvents the need for computational reconstruction. This approach offers a distinct advantage for accurately identifying and quantifying alternative splicing events, particularly for distinguishing between structurally similar isoforms, thus revealing transcript diversity with high accuracy and depth (Stark, Grzelak, and Hadfield 2019).

Our results show that hypoxia significantly altered alternative splicing (AS) events in the gills of *M. salmoides*. Exon skipping (SE) was the predominant change, marked by decreased exon inclusion, while alternative 3’ splice site (A3SS) events showed increased inclusion of variable regions. SE events, by eliminating one or more exons from mature mRNA, may result in the deletion or alteration of specific structural domains of proteins, thereby affecting their conformation, function, stability, or sites of interaction with other molecules (Nilsen and Graveley 2010). In conditions of hypoxia, this selective jumping of exons may be considered a potential adaptive strategy that enables cells to rapidly produce protein isoforms with modified functionality better suited to a low-oxygen environment (e.g., altered protein activity or reduced translational energy expenditure) (Sena et al. 2014; Ren et al. 2022). Indeed, it has been demonstrated that hypoxia can lead to an increase in global exon skipping (SE) (de Oliveira Freitas Machado et al. 2023). Furthermore, the selection of A3SS events results in alterations to the 5’ boundaries of downstream exons, which in turn may have consequences for the C-terminal region of the protein. This region may be important for subcellular localization, stability, or the regulation of enzymatic activity of the protein (Black 2003). A3SS events under hypoxia induction tend to increase the inclusion of variable regions, suggesting that the specific protein isoforms produced may have gain-of-function or specific regulatory roles in hypoxia adaptation. Notably, AS provides a level of regulation that is often independent of differential gene expression, providing additional fine-grained regulatory mechanisms for organisms to cope with environmental stress (Cao et al. 2025). The specific hypoxia-driven AS events and their resulting transcript variants revealed by PacBio Iso-Seq not only deepen our understanding of the molecular mechanisms of hypoxia adaptation in *M. salmoides*, but also highlight the importance of comprehensively resolving transcript diversity using long-read sequencing technologies when studying stress response. These findings establish a foundation for subsequent investigation of the specific functions of particular transcript isoforms in hypoxia tolerance.

### 4.2 Signaling pathways associated with hypoxia

When fish are exposed to low oxygen, cells induce a hypoxic response to adapt to the environment. The hypoxic response activates various signaling cascades that coordinate respiration adjustment, metabolism, and cell survival (Nakayama 2009). Our findings revealed pathways associated with the hypoxia response. For example, the “hypoxia inducible factor 1 subunit alpha (*hif1al*)” and “hypoxia inducible factor 1 subunit alpha inhibitor (*hif1an*)” genes were significantly upregulated with fold changes of 2.05 and 2.49 times in the hypoxic state compared to the normoxic condition, and they were significantly enriched in the GO term “cellular response to decreased oxygen levels”, “cellular response to hypoxia”, and “response to decreased oxygen levels” (Table S5), suggesting the involvement of the HIF-1 pathway plays a critical role in the hypoxic response of *M. salmoides*, which is consistent with other study that HIF-1 and mTOR signaling pathways play a critical role under hypoxic stress (Majmundar, Wong, and Simon 2010).

Hypoxia-inducible factor-1 (HIF-1) serves as a key transcription factor responsible for the cellular response to hypoxic environment, and mediating the expression of numerous genes involved in angiogenesis, cellular metabolism, and survival (Xiao 2015b; Kaelin and Ratcliffe 2008; Filice et al. 2022). Under normoxic conditions, HIF1α activity is suppressed by Hif1an, which hydroxylates conserved asparagine residues in HIF1α, hindering coactivator recruitment. Under hypoxic conditions, HIF1α stabilizes and dimerizes with HIF1β to form active HIF-1 (Long et al. 2015). Notably, in Gilthead seabream (*Sparus aurata*), significant upregulation of hif1an and hif1al genes in the HIF-1 signaling pathway has been observed during hypoxia exposure, supporting their role in activating HIF-1α (Raposo de Magalhaes et al. 2024).

### 4.3 Hypoxia impaired immune response in M. salmoides gill tissue

Hypoxia-induced signaling significantly impacts immune gene regulation, as evidenced by the enrichment of “Lipid and atherosclerosis” pathways, which are linked to white blood cell adhesion and transendothelial migration, suggesting hypoxia-mediated innate immune activation and inflammatory initiation (Jie et al. 2011). The TNF signaling pathway plays an important role in a variety of physiological and pathological processes, including cell proliferation, apoptosis, and inflammation induction (Aoki et al. 2008; Loo and Bertrand 2022). In this study, Tumor factor alpha (TNF-α), a key pro-inflammatory cytokine, was significantly downregulated across multiple immune-related pathways, such as Cytokine-cytokine receptor interaction, Toll-like receptor signaling, Apoptosis, and RIG-I-like receptor signaling, under hypoxic conditions (Sethi and Hotamisligil 2021; Zhong, Carvalho, et al. 2023). This downregulation suggests that *M. salmoides* may employ an immune-regulatory or suppressive strategy to mitigate excessive inflammatory responses during hypoxia.

Cytokines are key mediators of the immune communication and host defense, with chemokines facilitating leukocyte migration and immune responses (O’Shea, Gadina, and Siegel 2019; Metzemaekers et al. 2016; Cai et al. 2024). As mucosa-associated lymphoid tissues (MALTs), gills harbor diverse leukocyte populations that mediate local immunity (PRESS and EVENSEN 1999; Zhong, Liu, et al. 2023). In our study, we found that the activities of immunity-related chemokines (e.g., *ccl4*, *ccl5*, *cxcl8*, and *cxcl10*) were all down-regulated in hypoxia, and the chemokine receptor binding activities were also reduced, consistent with the response observed in large yellow croaker (*Larimichthys crocea*) under hypoxic stress (Mu, Li, Wei, et al. 2020). Hypoxia may inhibit the expression of immune-related genes by inducing a stress response, leading to reduced chemokine production. This could be due to a reduced energy supply, resulting in the preferential inhibition of non-essential functions, including immune signaling. The down-regulation of chemokines may decrease leukocyte recruitment and migration, thereby impairing local immune responses. Addtionally, the microenvironment of the gills may be significantly altered under hypoxic conditions. Such changes may affect cytokine stability, secretion patterns, and binding efficiency to receptors, further weakening the immune response. However, this suppression of immune function may be partially compensated by other pathways or alternative immune mechanisms. In contrast, the genes related to “oxygen binding” and “oxidoreductase activity” were up-regulated and significantly enriched under hypoxic conditions (Table S4). This suggests that hypoxic stress may alleviate oxygen deficiency by regulating metabolic processes and enhancing O_2_ uptake. More importantly, this suggests a strategic shift in energy utilization, prioritizing essential metabolic function over immune response under hypoxia.

The present study demonstrated that the hypoxic response is closely linked to immune response, and the repression of multiple chemokine-mediated signaling pathways under hypoxia may represent a resource-conserving response to oxygen limitation. However, this adaptive strategy may weaken the local immune defenses and potentially increase the risk of pathogen infection in fish.

### 4.4 Hypoxia inhibited cell proliferation in M. salmoides

The genes *rpa1*, *mcm5*, *mcm2*, *dna2*, *mcm4*, *mcm6* and *prim1* were enriched in the DNA replication signaling pathway. The MCM complex, a replicative helicase essential for DNA replication, unwinds the template DNA and assembles replisomes (Pellegrini et al. 2020; Jenkyn-Bedford et al. 2021; Jones et al. 2021). MCM family genes are important marker genes in the regulation of DNA replication and cell proliferation (Davuluri et al. 2008). In addition, *mcm5*, *mcm2*, *mcm4*, *mcm6* and *mcm3* were also enriched in the cell cycle pathway, and all of these genes were down-regulated. Furthermore, the genes related to DNA replication and cell cycle, which were among the most significantly enriched KEGG pathways, were also down-regulated, suggesting that cell proliferation in *M. salmoides* gill tissues may be inhibited.

Differentially expressed genes in Salmonids and *Takifugu rubripes* gill tissues under hypoxia were enriched in cell cycle and proliferation-related pathways, with altered expression patterns of growth-inhibitory genes (Akbarzadeh et al. 2020; Shang et al. 2022). Hypoxia has been shown to inhibit cell proliferation in a variety of cell types, and the reduction in cell number may reduce overall oxygen demand under hypoxic conditions (Hubbi and Semenza 2015). These hypoxia-repressed genes may reflect a physiological change that stops cell proliferation and translational activity, thereby conserving energy under metabolic stress (Chi et al. 2006). Similarly, genes involved in cell cycle progression were down-regulated in zebrafish embryos and in hepatocytes of hybrid striped bass (*Morone saxatilis* × *Morone chrysops*) under acute and chronic hypoxia (Ton, Stamatiou, and Liew 2003; Beck et al. 2016). Thus, the down-regulation of gill cell proliferation may represent a physiological strategy for *M. salmoides* to cope with hypoxic environments. Meanwhile, inhibition of cell growth and proliferation may help redirect important energy resources to metabolic processes that are essential for hypoxic survival, such as anaerobic metabolism and stress response pathways.

### 4.5 Hypoxia promoted apoptosis in M. salmoides cells

Apoptosis is a programmed cell death process essential for tissue development, metabolic homeostasis, and immune regulation. Caspases, key cysteine proteases, remain inactive as procaspases until activated by apoptotic signals, leading to cell death (Sahoo et al. 2023). Caspase 7 was up-regulated in KEGG-significantly enriched pathways, including Legionellosis, Apoptosis, Pertussis, Lipid and atherosclerosis, the TNF signaling pathway, and Non-alcoholic fatty liver disease, all with significant enrichment (P < 0.05). Once activated, Caspase 7 clears and activates various key intracellular proteins, including structural proteins, DNA repair proteins, and other effector caspases. Caspase 7 activation leads to nuclear fragmentation, cytoplasmic shrinkage, chromatin condensation, and membrane disruption, which are typical features of apoptosis (Van Opdenbosch and Lamkanfi 2019). This apoptotic process is an important mechanism for maintaining tissue homeostasis and inhibiting the proliferation of abnormal cells. Similar apoptosis responses have been reported in *M. salmoides* and hybrid yellow catfish (*Tachysurus fulvidraco*♀×*Pseudobagrus vachellii*♂) (Sun et al. 2020; Tao et al. 2021). Therefore, hypoxic stress induces apoptosis by triggering the activation of apoptosis-related genes in response to cell membrane and protein damage, thereby maintaining the stability of the body’s internal environment by removing damaged cells.

## 5. Conclusion

In summary, the present study investigated the histological and molecular biological changes in the gills of *M. salmoides* under hypoxic stress. We provided a high-quality transcriptome for *M. salmoides* by integrating full-length transcriptome sequencing (Iso-Seq) and high-throughput RNA-Seq, offering valuable genetic resources for future studies. Histological analysis revealed adaptive changes and apparent structural alterations in gill tissues under hypoxic conditions, while comparative transcriptomic analyses indicated that most DEGs were down-regulated, particularly those involved in energy metabolism, immune response, and morphological regulation. GO and KEGG analysis indicated that hypoxia-induced DEGs were enriched in pathways related to energy metabolism, immune response, cell growth and proliferation, apoptosis, and chemokine signaling, all of which may help limit hypoxia-induced damage. The results of this study not only enhance our understanding of the regulatory mechanism underlying the hypoxic response in *M. salmoides*, but also provide valuable insights to help prevent and reduce production loss in aquaculture.

## Acknowledgments

This study was supported by the APRC-CityU New Research Initiatives/Infrastructure Support from Central (9610574, 7006064) from City University of Hong Kong.

## CRediT authorship contribution statement

**Xiaoli Ma:** Conceptualization, Resources, Methodology, Investigation, Writing – review & editing. **Baofeng Su:** Writing – original draft preparation, Formal analysis. **Anxin Ye:** Writing – original draft preparation, Visualization, Investigation, Formal analysis. **Renyu Wan:** Investigation, Formal analysis, Conceptualization. **Keyan Zhao:** Visualization, Formal analysis, Data curation. **Liang Zhong:** Data curation, Formal analysis. **Wenxin Yu:** Data curation, Formal analysis. **Yuhe Liu:** Investigation, Formal analysis. **Jinyi Wu:** Data curation. **Wenlong Cai:** Resources, Funding acquisition, Conceptualization, Investigation, Writing – review & editing.

## Data availability

We have deposited all raw sequencing data to the National Center for Biotechnology Information (NCBI) under BioProject accession numbers PRJNA1498806 and PRJNA1501902.

## Appendix A. Supplementary data

Table S1 Statistics of sequence categories in the full-length transcriptome of *Micropterus salmoides* under hypoxia challenge

Table S2 Gene annotation results for Micropterus salmoides

Table S3 Summary of Illumina RNA-Seq data

Table S4 Differentially expressed genes (DEGs) from the Illumina RNA-Seq data

Table S5 GO enrichment results of EG vs. CG.

Table S6 KEGG enrichment results of EG vs. CG.

Table S7 Statistical analysis of gill filament length and width in hypoxic and normoxic groups

