## Supplementary material for "Integrated Transcriptomics and Histological Analysis Reveals Gill Adaptation Mechanisms to Hypoxic Stress in Largemouth Bass (*Micropterus salmoides*)": Figure S1 and Figure S2

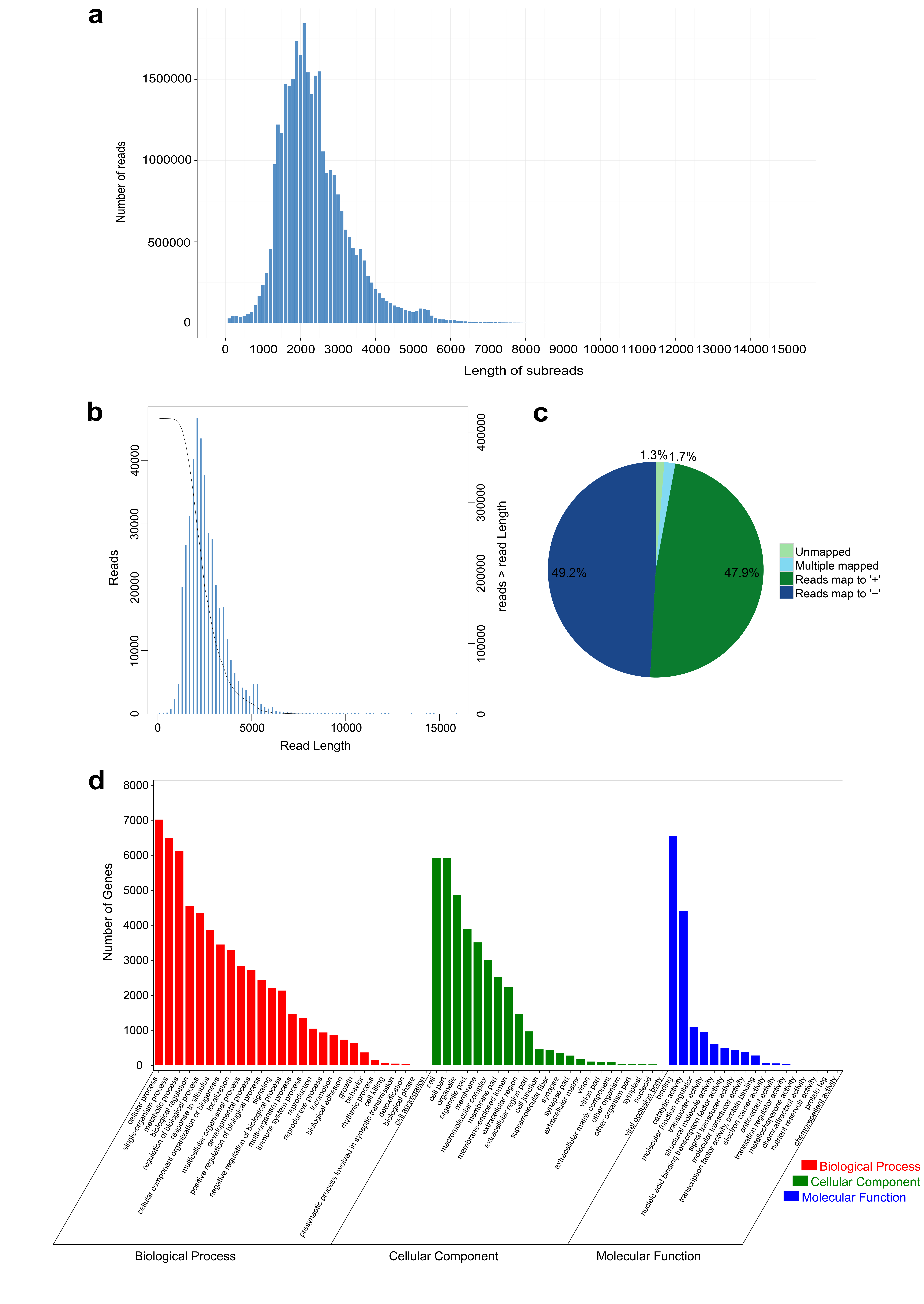
Fig. S1 (a) Length distribution of subreads. The X-axis represents the length of subreads, and the Y-axis represents the number of subreads. (b) Length distribution of FLNC (full-length non-chimeric) reads. The X-axis represents the length of FLNC reads. The left Y-axis (histogram) indicates the number of reads within a specific length range (X-axis), while the right Y-axis indicates the cumulative number of reads with lengths greater than a specific value (X-axis). (c) Alignment results of high quality (HQ) isoforms with the reference genome. (d) GO enrichment results from full-length transcriptome sequencing, showing three main categories: biological processes, cellular components, and molecular functions.

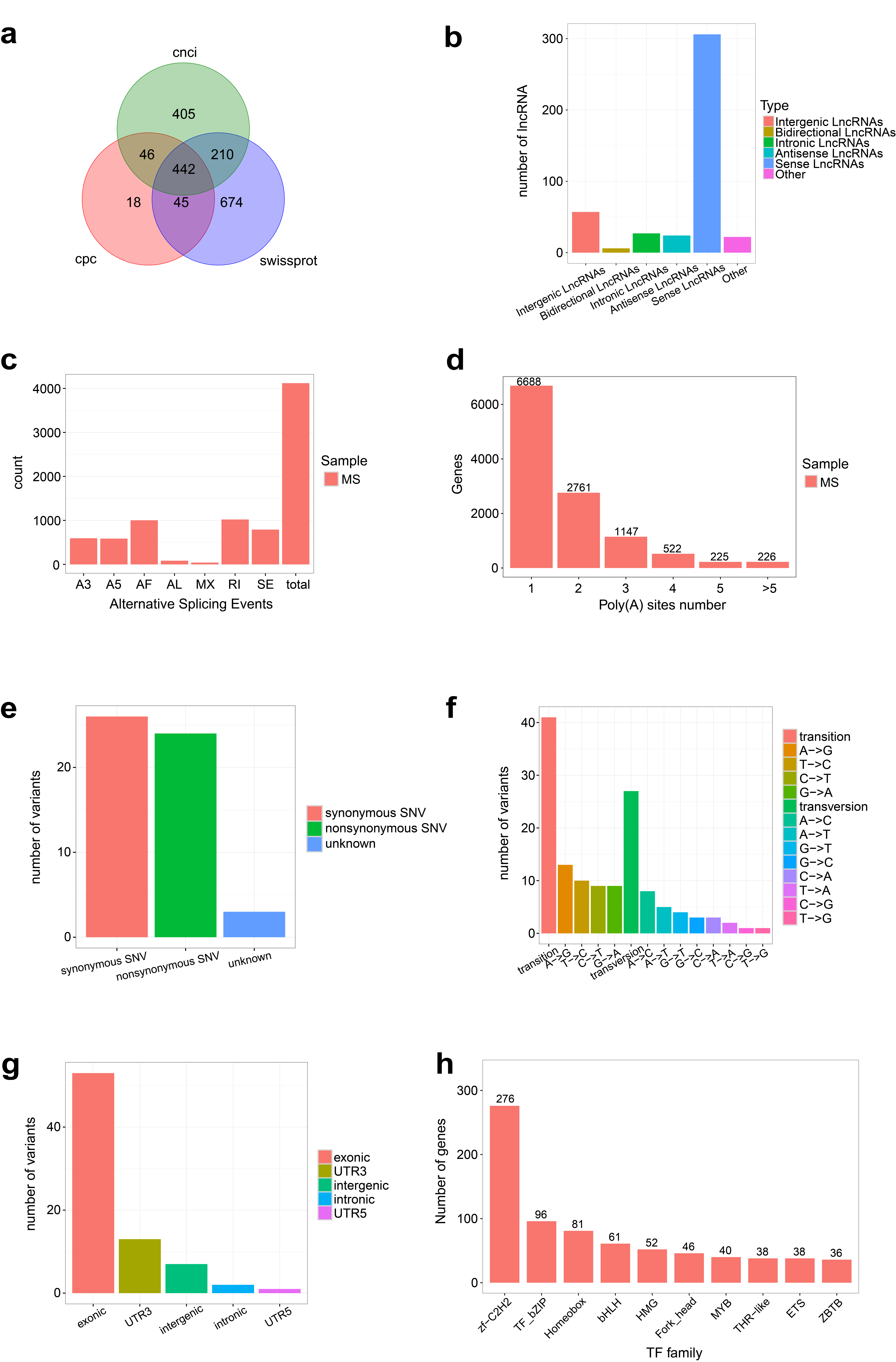

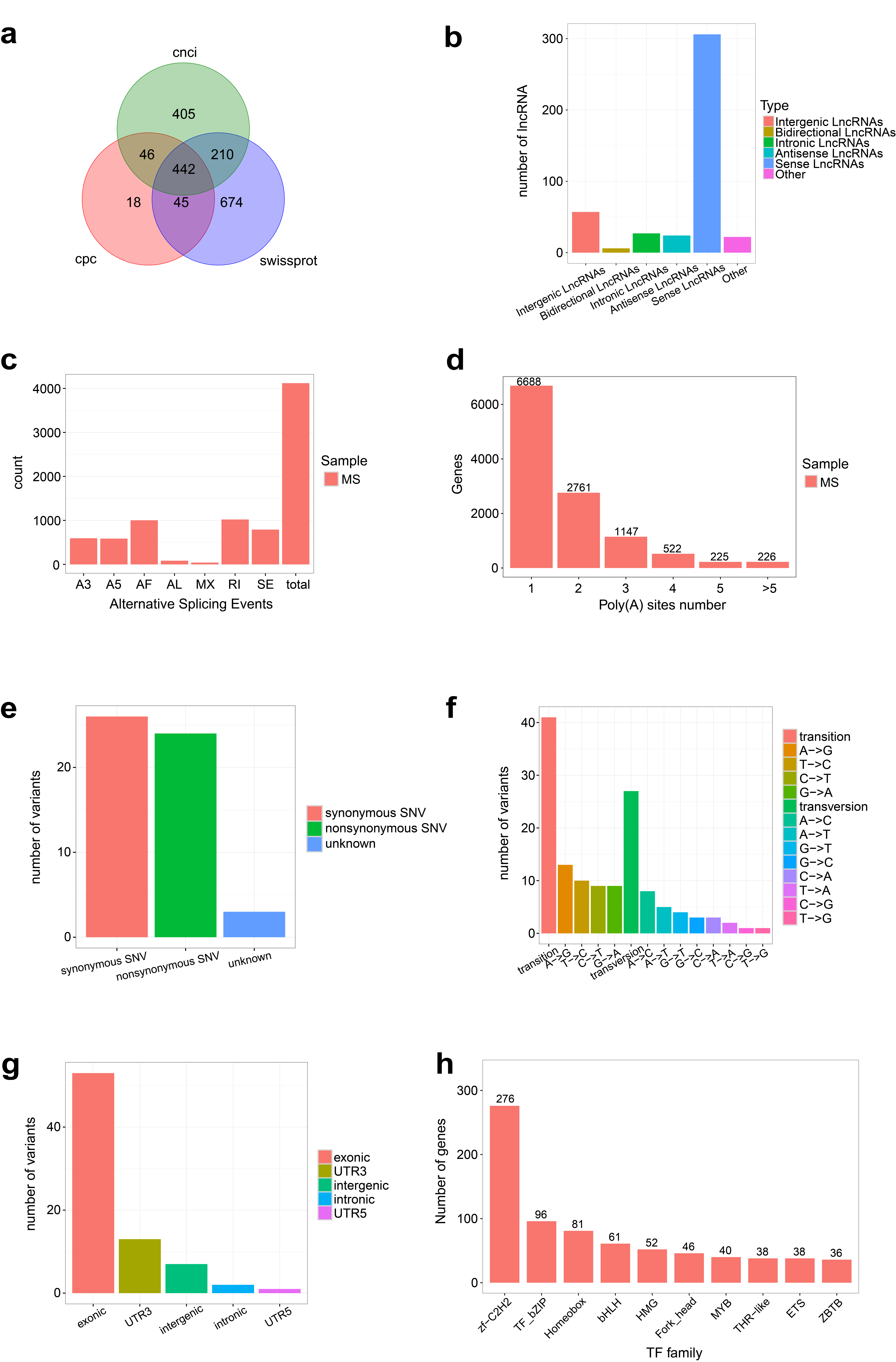

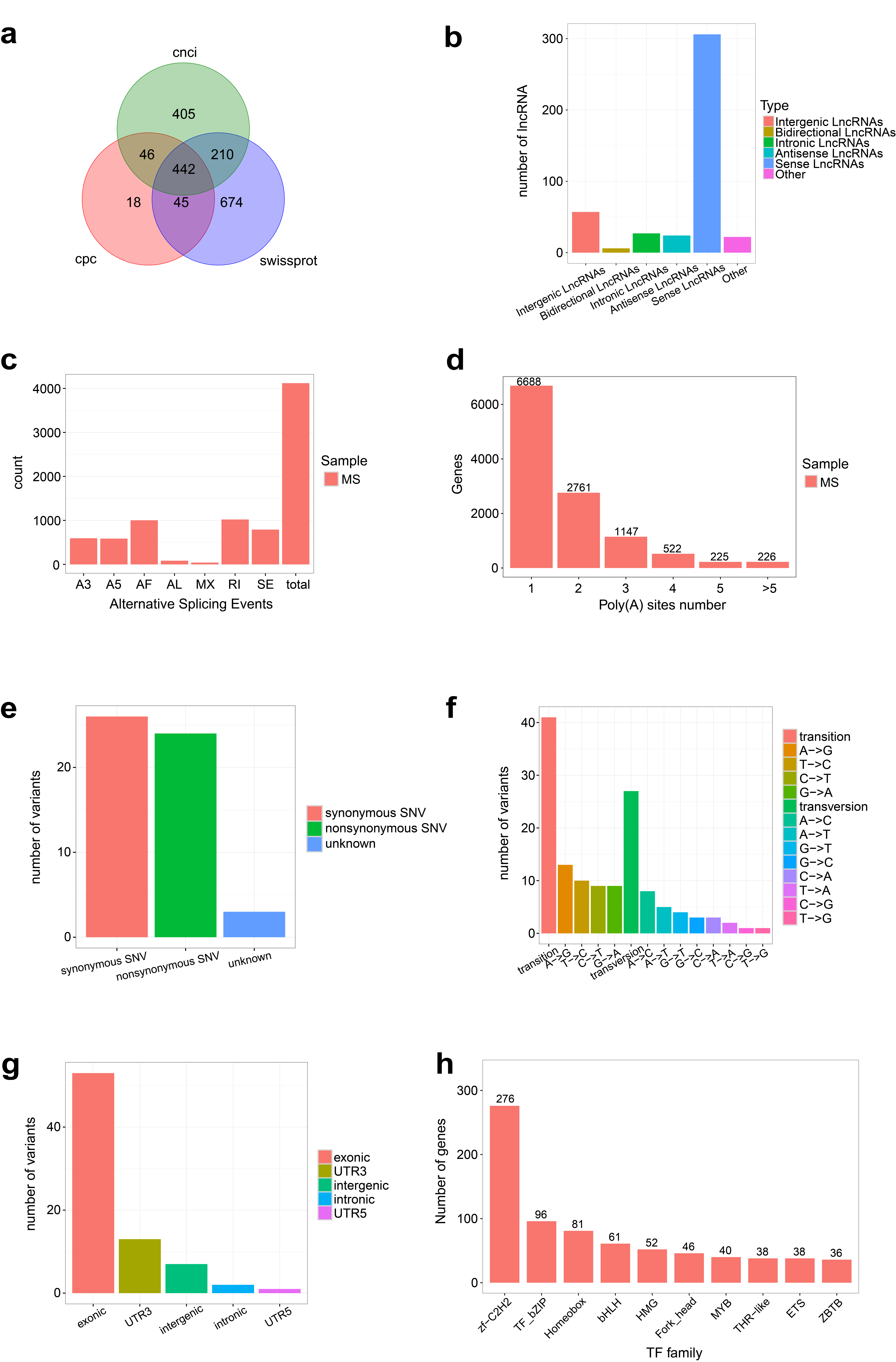

**a**

**b**

**c**

**d**

Fig. S2 (a) Venn diagram of lncRNA prediction results based on CNCI, CPC, and Swiss-Prot databases. (b) Summary of lncRNA classification results. (b) Polyadenylation profile. The X-axis represents the number of polyadenylation (polyA) sites, and the Y-axis represents the number of genes. (c) Functional classification of SNPs. The X-axis represents the classification of variants (synonymous SNV and nonsynonymous SNV), and the Y-axis represents the number of variants. (d) Summary of SNP mutation types.
